# A *de novo* reference genome of the golden jackal, *Canis aureus*

**DOI:** 10.1101/2025.10.09.681322

**Authors:** Sven Winter, René Meißner, Stefan Prost, Carola Greve, Charlotte Gerheim, Marine Arakelyan, Sargis Aghayan, Astghik Ghazaryan, Jennifer Hatlauf, Pamela A. Burger

## Abstract

The golden jackal (*Canis aureus*) is rapidly expanding its range in Europe, driven by climate and habitat changes, human influence, and changes in competition with wolves. Its ecological flexibility enables it to thrive in various habitats, including urban areas, raising concerns about its potential role in spreading zoonotic diseases. Jackals may act as reservoirs for pathogens such as Lyme disease and babesiosis, affecting wildlife, humans, and pets. Their close genetic relationship with domestic dogs also increases the risk of hybridization and host-jumping, complicating disease dynamics.

To better understand their dispersal ability and host-pathogen dynamics, we present the first chromosome-level genome assembly of the golden jackal, generated using PacBio HiFi sequencing and reference-based scaffolding. The final assembly has a total length of 2.53 Gb in 325 scaffolds, with 98.41% of the sequence anchored to the expected 38+XY chromosomes. The assembly shows high contiguity, with scaffold and contig N50 values of 68.03 Mb and 56.64 Mb, respectively. Annotation revealed 26,084 protein-coding genes, and repetitive elements account for 40.58% of the total assembly.

This high-quality reference genome provides an essential resource for studying the genetic basis of the golden jackal’s adaptation, ecological interactions, and potential as a zoonotic reservoir. It also supports efforts to monitor population expansion and its effects on ecosystems. By advancing our understanding of golden jackal genetics, this work enables future research on evolution, host-pathogen dynamics, and the broader consequences of wildlife dispersal in a rapidly changing environment.

## Introduction

Canids (*Canidae*) are a diverse group of mostly carnivorous mammals that exhibit a wide range of ecological adaptations and social behaviors (Allen, 2017). The approximately 35 species are distributed across all continents except Antarctica, highlighting their ability to thrive in vastly different environments (Krofel et al., 2022). One such species, the golden jackal (*Canis aureus* LINNAEUS, 1758), is a mid-sized canid that has been experiencing an accelerated natural dispersal, particularly within Western Asia and into Europe (Hatlauf et al., 2021). This expansion is thought to be driven by a combination of several factors, including climate change, shifts in habitat, land-use changes, and other human activities, all of which have facilitated its range expansion (Krofel et al., 2017; Spassov & Acosta-Pankov, 2019).

The golden jackal’s notable adaptability allows it to occupy a wide array of habitats, ranging from forests and grasslands to urban environments (Salek et al., 2014; Torretta et al., 2020). Its opportunistic feeding habits, which include a broad diet of small mammals, birds, insects, fruits, and vegetables, make it highly resilient and capable of colonizing new areas (Lanszki et al., 2022). While the presence of golden jackals in new ecosystems can provide benefits, such as natural pest control by regulating populations of rodents and other small vertebrates (Lanszki et al., 2016), the species’ expansion also presents potential challenges (Gherman & Mihalca, 2017). As jackals move into new areas, they may compete with native species for resources and potentially disrupt local ecosystems (Krofel et al., 2017; Serva et al., 2023). Additionally, the golden jackal’s capacity to survive in urban environments brings it into closer proximity with humans, which increases the likelihood of interaction with domestic animals, raising concerns about zoonotic disease transmission (Hatlauf, 2024; Lapid et al., 2023).

Recent studies have focused on the potential public health risks associated with the movement of golden jackals, especially in light of the discovery of jackals in Denmark carrying tick vectors and zoonotic pathogens (Klitgaard et al., 2017). Pathogens, such as those transmitted by ticks, can pose significant risks to both wildlife and human health, which underscores the importance of monitoring the spread of this species. The potential for golden jackals to carry and transmit diseases like Lyme disease and babesiosis demonstrates how their expanding range could influence the epidemiology of zoonoses, particularly in regions where the jackal may interact with domestic animals and humans (Lymbery et al., 2014). Furthermore, the close phylogenetic relationship between golden jackals and domestic dogs (*Canis lupus familiaris*) raises the possibility of hybridization and host jumping, which could exacerbate the spread of zoonotic diseases or introduce novel pathogens into local ecosystems (Galov et al., 2015; Gherman & Mihalca, 2017; Stefanovic et al., 2024).

Given the important ecological role golden jackals play in their current and future habitats, they serve as important models for understanding climate-mediated dispersal of zoonotic diseases. Their expansion is a good example of how environmental changes and human-induced factors influence the movement of wildlife and the pathogens they carry. Studying jackals offers valuable insights into the broader dynamics of host-pathogen coevolution and the transmission of diseases across species boundaries. To support these efforts, a comprehensive genomic resource is essential.

In this study, we present the first chromosome-level genome assembly of the golden jackal, generated using PacBio HiFi long-read sequencing, supplemented by reference-based scaffolding. This high-quality reference genome provides an invaluable tool for investigating the genetic basis of the species’ adaptation to new environments and its interactions with both its ecosystem and pathogens. It also enables more effective monitoring of the species and the diseases it may carry, facilitating a better understanding of how climate change and landscape fragmentation contribute to the spread of zoonotic diseases. By advancing our genetic knowledge of golden jackals, we provide a foundation for future studies on the evolution, population dynamics, and ecological impacts of this increasingly prominent species.

## Materials and Methods

### Sampling and Sequencing

Tissue samples from a subadult male golden jackal (sample ID: CA_001; PathoNo.: SK/123/23) shot by a hunter in Villach, Carinthia, Austria, were collected during post mortem examination at the University of Veterinary Medicine Vienna, Austria, on 06.02.2023 and stored frozen at −80°C. High molecular weight DNA was extracted from spleen tissue using the innuPrep SE HMW kit (IST Innuscreen GmbH, Germany) according to the manufacturer’s recommendations. The quantity and quality of the DNA were evaluated using the QuantiFluor ONE dsDNA System on the Quantus Fluorometer (Promega Corporation, USA) and the Genomic DNA Screen Tape on the Agilent 4150 TapeStation system (Agilent Technologies, Inc., USA), respectively.

Library preparation for Pacific Biosciences of California, Inc (PacBio) circular consensus sequencing (CCS) was prepared using the SMRTbell Express Prep Kit 3.0 (PacBio, USA) and sequenced on the PacBio Revio system.

### Assembly and QC

The PacBio-generated raw data BAM file was converted into FASTQ format using BAM2fastx v.1.3.0, a PacBio Secondary Analysis Tool (https://github.com/PacificBiosciences/pbbioconda). The initial assemblies, including two, which resemble the two potential parental haplotypes, were generated with hifiasm v.0.24.0 (RRID:SCR_021069) (Cheng et al., 2021) using standard parameters. As sample quality was insufficient for a proximity-ligation library, we did not aim for a haplotype-resolved assembly but instead continued with the pseudohaploid assembly for the next steps. As we were unable to use Hi-C data for the scaffolding, due to the fact that the sample was stored in ethanol, we instead used the Mexican gray wolf assembly (Accession No. GCF_048164855.1) for a reference-based scaffolding with RagTag v.2.1.0 (Alonge et al., 2022). Subsequently, gaps in the scaffolded assembly were breached using the original PacBio long-read data with TGS_gapcloser v.1.2.1 (RRID:SCR_017633)(Xu et al., 2020). The mitochondrial genome was assembled using the MitoHiFi workflow (RRID:SCR_026369) in Galaxy v.3.2.3 (RRID:SCR_006281)(Blankenberg et al., 2014; Uliano-Silva et al., 2023).

### Assembly Quality Assessment

Assembly statistics to evaluate contiguity of the assembly were generated with Quast v.5.0.2 (<u>RRID:SCR_001228</u>)(Gurevich et al., 2013). The completeness of the assembly was assessed by gene set completeness analyses using BUSCO v.5.8.3 (<u>RRID:SCR_015008</u>) (Manni et al., 2021) and compleasm v.0.2.7 (<u>RRID:SCR_026370</u>)(Huang & Li, 2023) against the carnivora_odb12 orthologs dataset, as well as based on k-mers (k=21) using Merqury v.1.3 with Meryl v. 1.3 (<u>RRID:SCR_022964</u>)(Rhie et al., 2021). Minimap2 v.2.28 (<u>RRID:SCR_018550</u>)(Li, 2018) and samtools v.1.21 (<u>RRID:SCR_002105</u>)(Danecek et al., 2021) were used to map the PacBio reads to the final assembly and sort the mapping file to check the mapping rate with Qualimap bamqc v.2.3 (<u>RRID:SCR_001209</u>)(Okonechnikov et al., 2016). The mapping file was also used together with the results from a Diamond v.2.1.10 (<u>RRID:SCR_016071</u>)(Buchfink et al., 2021) search against Swiss-Prot (release 2025-01) (The UniProt Consortium, 2019) in Blobtools v. 1.1 (<u>RRID:SCR_017618</u>)(Laetsch & Blaxter, 2017) to check for potential contamination.

### Annotation

Repeats in the assembly were annotated with RepeatMasker v.4.1.0 (<u>RRID:SCR_012954</u>) (Smit et al., 2015) using a custom *de novo* repeat library generated from the presented assembly with RepeatModeler v.2.0.1 (<u>RRID:SCR_015027</u>) (Flynn et al., 2020) combined with the Canidae specific repeats from Dfam 3.1 and RepBase (release 20181026) (Bao et al., 2015; Storer et al., 2021). Interspersed repeats were hardmasked (Ns) while simple repeats were softmasked (lowercase) in preparation for the gene annotation.

Genes were predicted using the homology-based approach implemented in the GeMoMa pipeline v.1.7.1 (<u>RRID:SCR_017646</u>) (Keilwagen et al., 2016, 2018) using MMseqs2 v. 13.45111 (<u>RRID:SCR_022962</u>) (Steinegger & Söding, 2017) as an alignment tool. The following six mammalian genomes and corresponding annotations were used as references: *Homo sapiens* (GCF_000001405.40), *Mus musculus* (GCF_000001635.27), *Canis lupus familiaris* (German Shepherd, GCF_011100685.1; Wang et al., 2021), *Vulpes vulpes* (GCF_048418805.1), *Vulpes lagopus* (GCF_018345385.1;, *Canis lupus baileyi* (GCF_048164855.1; Rhie et al., 2021).

Subsequently, the proteins predicted by GeMoMa were functionally annotated by a BLASTP v.2.11.0+ (<u>RRID:SCR_001010</u>) (Camacho et al., 2009) search against the Swiss-Prot database (<u>RRID:SCR_002380; release 2025-01</u>) (The UniProt Consortium, 2019) with an e-value cutoff of 10^-6^. Gene ontology (GO) terms, domains, and motifs were further annotated using InterProScan v.5.59.91 (<u>RRID:SCR_005829</u>) (Jones et al., 2014; Quevillon et al., 2005). BUSCO v.5.8.3 (<u>RRID:SCR_015008</u>) (Manni et al., 2021) and compleasm v.0.2.7 (<u>RRID:SCR_026370</u>) (Huang & Li, 2023) were used to evaluate gene completeness in reference to the carnivora_odb12 dataset.

## Results and Discussion

Sequencing of the two prepared CSS libraries on the PacBio Revio platform resulted in 6,520,091 HiFi reads (86.95 Gb) with a mean read length of 13.34 kb and a mean quality of 29.7 (Supplementary Material S1). The initial assembly with Hifiasm resulted in a draft assembly of 2.54 Gb with 502 contigs and a contig N50 of 55.91 Mb (Table 2A). Due to the sample quality, we were unable to generate proximity ligation libraries for scaffolding and opted for a reference-based scaffolding approach. After removing some of the smallest contigs, due to potential bacterial contamination (taxonomic assignments, variation in sequencing depth and GC-content) identified by BlobTools (Fig. 2C; Supplementary Material S2), the final highly contiguous chromosome-level assembly has a total length of 2.53 Gb with 325 scaffolds and a scaffold and contig N50 of 68.03 MB and 56.64 Mb, respectively (Table 2A). Of the 325 scaffolds (incl. the mitochondrial genome), the largest 40 (> 7 Mb) corresponding to the expected 38 autosomes and the two allosomes (Tanomtong et al., 2015) span 98.41% of the total assembly length, resulting in a L50 of 15 scaffolds. Of the 13,727 BUSCO genes in the carnivora_odb12 dataset, BUSCO identified 13,392 (97.6%) as complete single copy and 158 (1.2%) as duplicated BUSCOs, while compleasm found 13,528 (98.55%) single copy and 126 (0.16%) duplicated BUSCOs (Fig. 2A). The high BUSCO and compleasm scores, as well as a high mapping rate of 99.88% and a mapping quality of 39.7, highlight the quality of the presented assembly. Furthermore, Merqury estimated a k-mer completeness score of 96.93% with a QV score of 63.57, corresponding to an error rate of 4.39e-07 (see also Fig. 2 B).

**Table 1.**
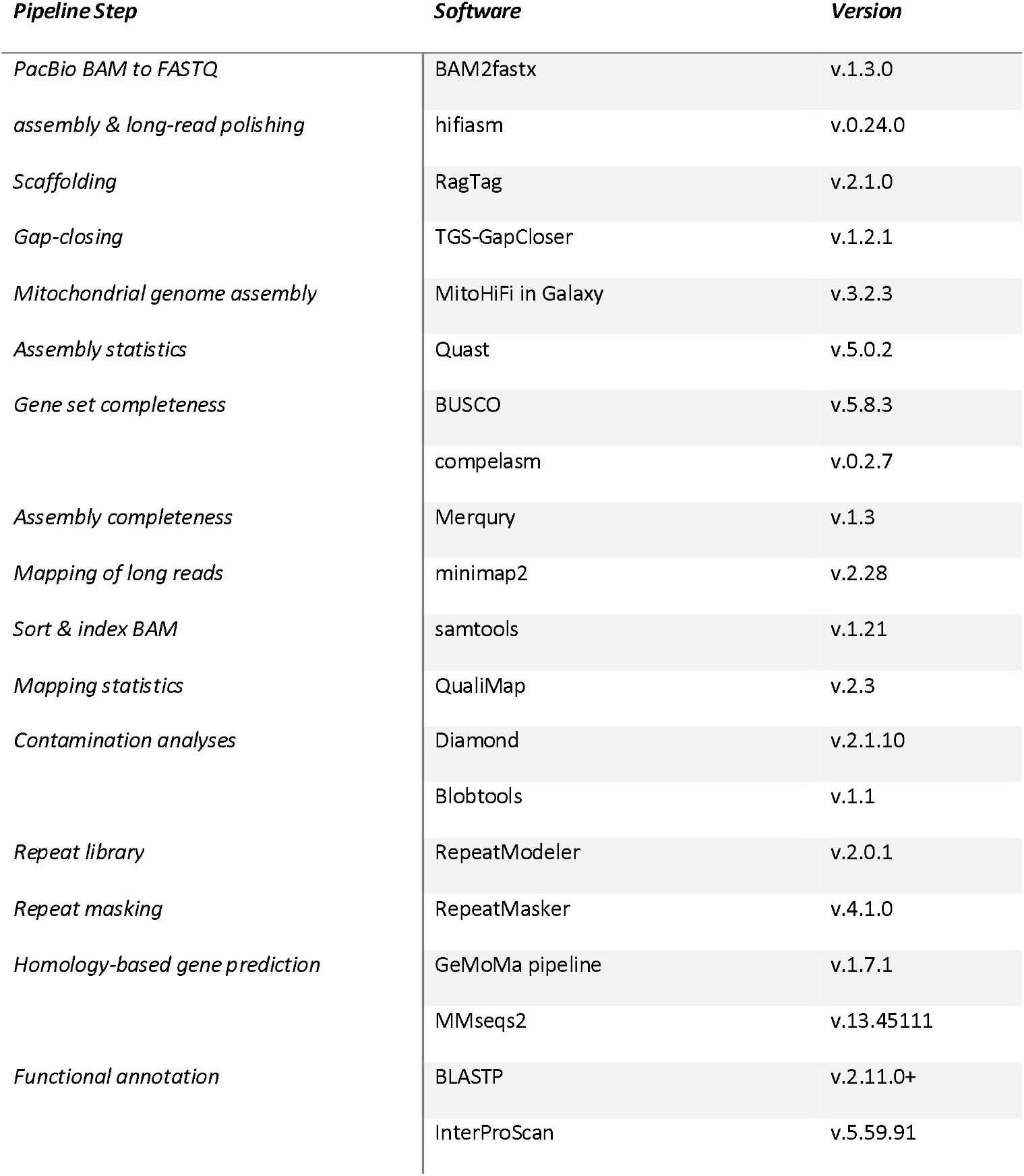
Software and versions used to generate the *Canis aureus* assembly.

**Table 2.**
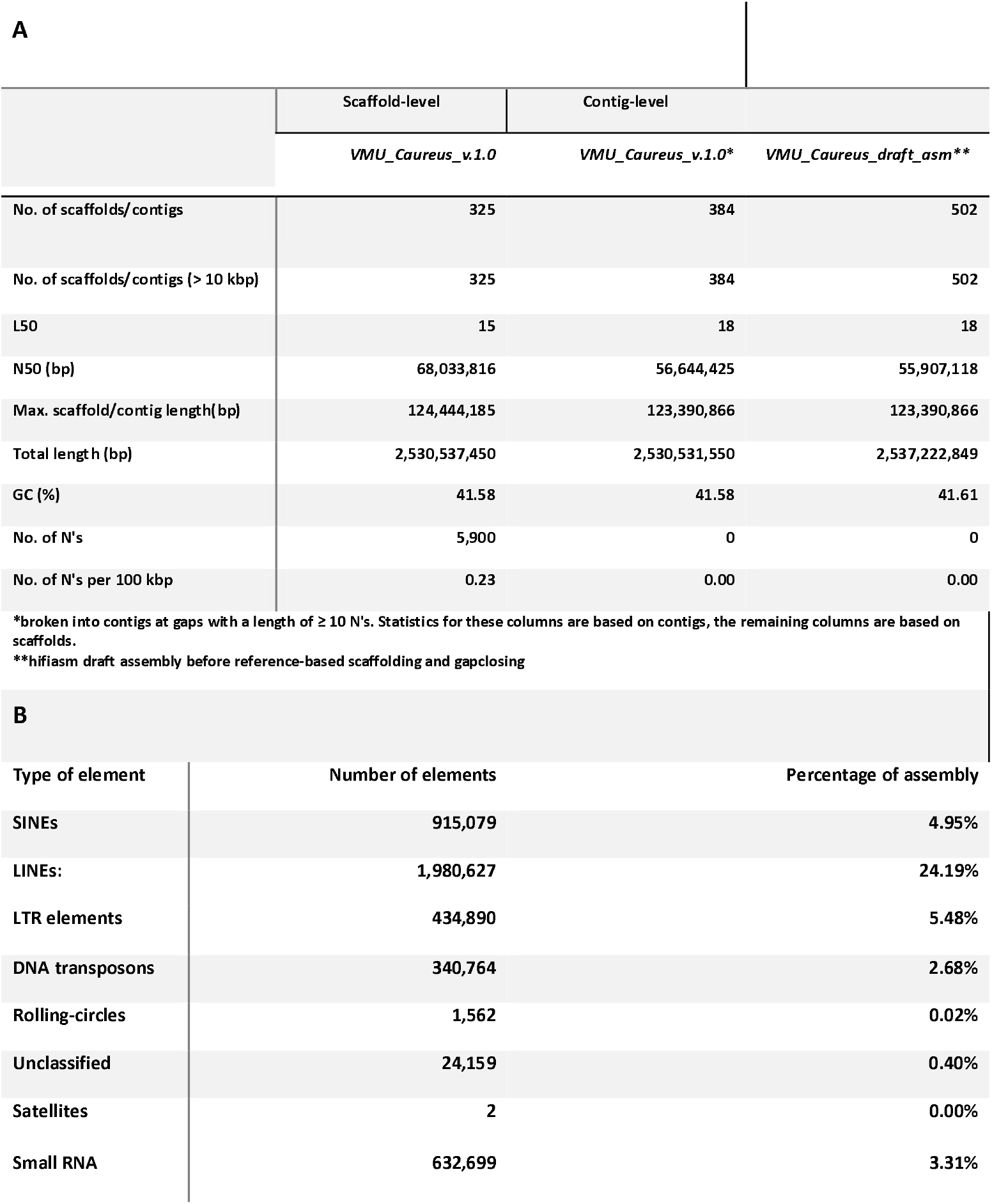
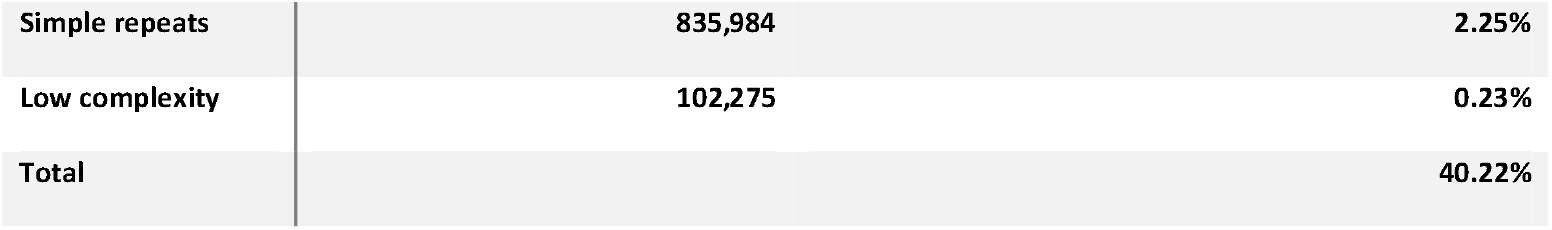
Assembly statistics of *VMU_Caureus_v.1.0* (A) and its repeat content (B)

| **A** |  |  |  |
| --- | --- | --- | --- |
|  | Scaffold-level | Contig-level |  |
|  | *VMU_Caureus_v.1.0* | *VMU_Caureus_v.1.0\** | *VMU_Caureus_draft_asm\*\** |
| No. of scaffolds/contigs | 325 | 384 | 502 |
| No. of scaffolds/contigs (> 10 kbp) | 325 | 384 | 502 |
| L50 | 15 | 18 | 18 |
| N50 (bp) | 68,033,816 | 56,644,425 | 55,907,118 |
| Max. scaffold/contig length(bp) | 124,444,185 | 123,390,866 | 123,390,866 |
| Total length (bp) | 2,530,537,450 | 2,530,531,550 | 2,537,222,849 |
| GC (%) | 41.58 | 41.58 | 41.61 |
| No. of N's | 5,900 | 0 | 0 |
| No. of N's per 100 kbp | 0.23 | 0.00 | 0.00 |
| *broken into contigs at gaps with a length of $\geq 10$ N's. Statistics for these columns are based on contigs, the remaining columns are based on scaffolds. | | | |
| **hifiasm draft assembly before reference-based scaffolding and gapclosing |  |  |  |
| **B** |  |  |  |
| Type of element | Number of elements | Percentage of assembly |  |
| SINEs | 915,079 | 4.95% |  |
| LINES: | 1,980,627 | 24.19% |  |
| LTR elements | 434,890 | 5.48% |  |
| DNA transposons | 340,764 | 2.68% |  |
| Rolling-circles | 1,562 | 0.02% |  |
| Unclassified | 24,159 | 0.40% |  |
| Satellites | 2 | 0.00% |  |
| Small RNA | 632,699 | 3.31% |  |

**Figure 1.**
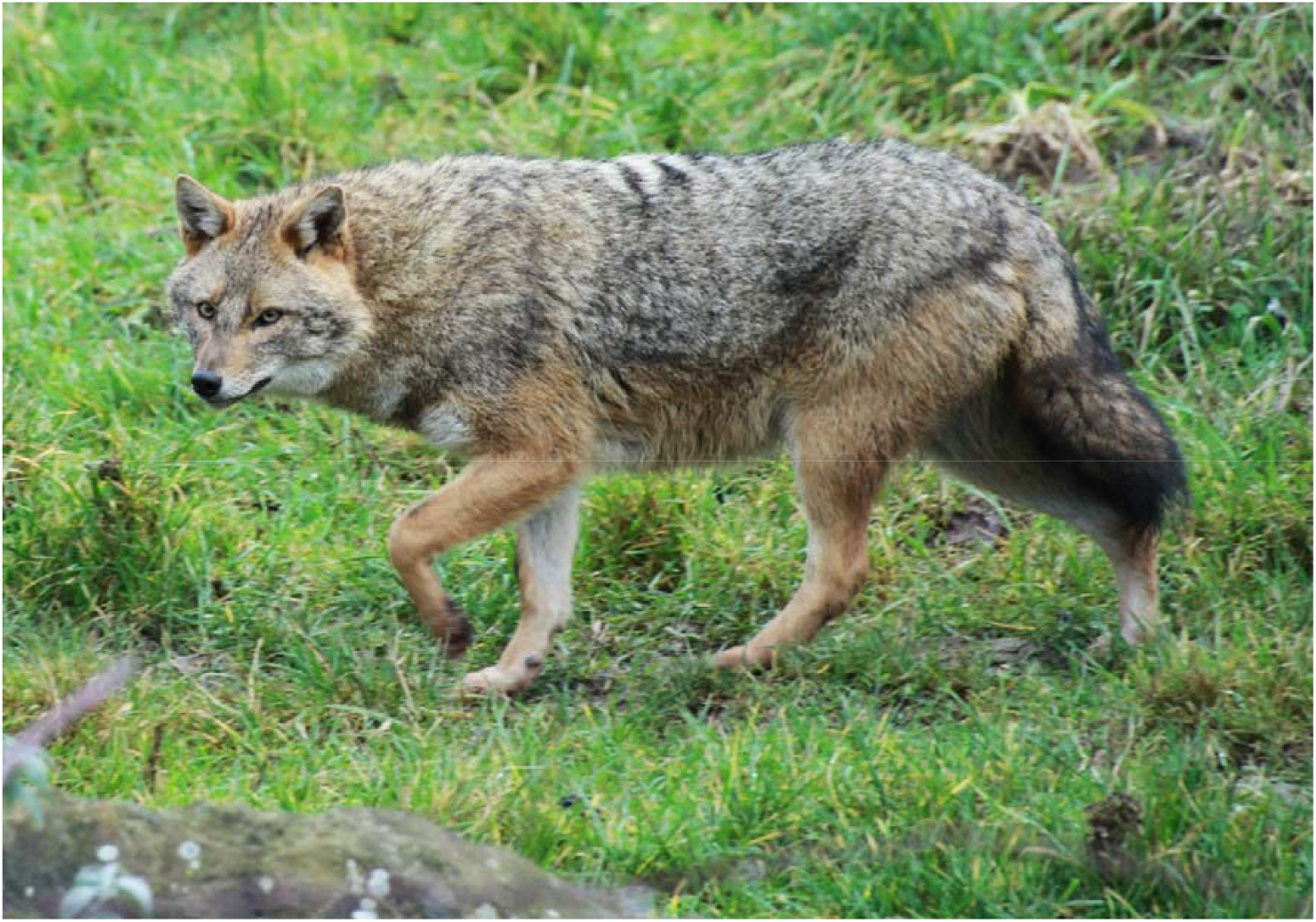
Golden jackal (*Canis aureus*) ©Jennifer Hatlauf.

**Figure 2.**
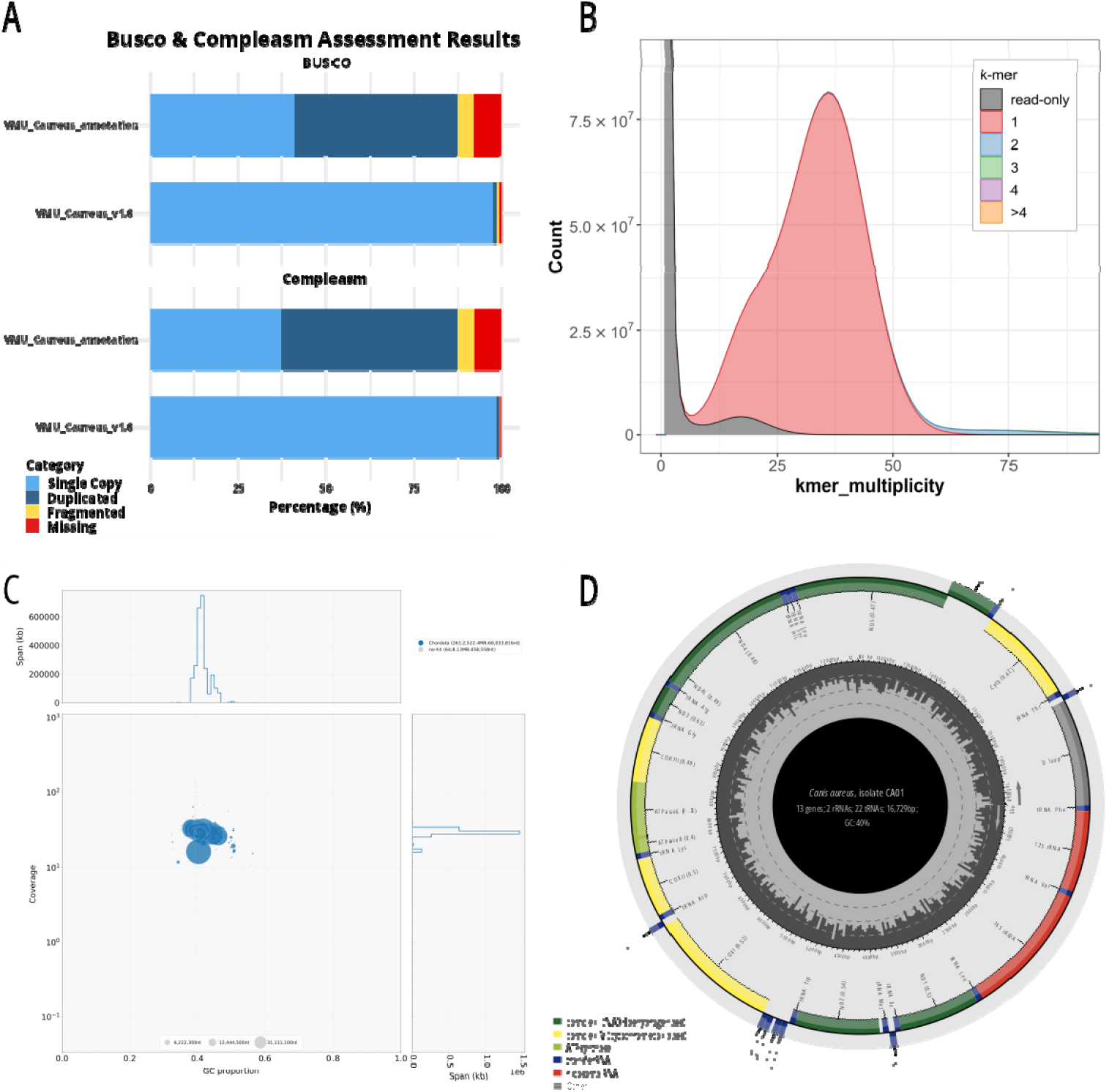
Assembly quality assessment and mitochondrial genome of *VMU_Caureus_v1.0*. (A) BUSCO and compleasm assessment results for the final assembly and the annotation using the carnivora_obd12 dataset. (B) Merqury k-mer copy number spectrum plot.(C) BlobPlot analysis comparing GC content (x-axis) and sequence coverage of the PacBio data (y-axis) after removal of potentially contaminated scaffolds, see Supplementary Material S2 for BlobPlot before removal. The color of the “blobs” represents taxonomic assignment of the scaffolds based on the Swiss-Prot database. (D) Annotation of the circular mitochondrial genome of CA01.

The mitochondrial genome assembled from the HiFi reads was circular with a length of 16,729 bp, following the standard composition and structure of vertebrate mitochondrial genome sequences with 13 protein-coding genes, 2 rRNAs, and 22 tRNAs (Fig. 2D).

### Annotation

#### Repeat annotation

Repeat annotation identified a repeat content in the final assembly *VMU_Caureus_v1.0* of 40.58% or 1.02 Gb (Table 2B). With 24.19% of the assembly Long Interspersed Nuclear Elements (LINEs) were the most common repeats, followed by LTR Elements (5.48%) and Short Interspersed Nuclear Elements (SINEs, 4.95%). Non-Retroelements such as DNA Transposons, small RNA, and simple repeats, each accounted for less than 4% of the assembly.

#### Gene annotation

Homology-based gene prediction with GeMoMA identified 26,084 genes in *VMU_Caureus_v.1.0*. The median gene length was 6,370 bp, spanning 469.26 Mb of the assembly. BUSCO identified 41.2% (n = 5,653) of the 13,727 Carnivora orthologous genes from the carnivora_odb12 BUSCO dataset in the assembly as single copy and 46.4% (n = 6,372) as duplicated, with 1,089 BUSCO genes (7.9%) missing (Fig. 2A). Compleasm identified 37.42% (n = 5,137) BUSCOs as single-copy and 50.11% (n = 6,879) as duplicated, with 7.71% (n = 1,058) missing (Fig. 2A). InterProScan functionally annotated 61,027 of the 61,813 predicted proteins (98.73%) and assigned 53,147 proteins (85.98%) to the Reactome and assigned at least one Gene Ontology (GO) term to 45,071 proteins (72.92%). Furthermore, 96.73 % (n=59,792) of the predicted proteins were matched to entries in the Swiss-Prot database.

## Conclusions

High-quality long-read-based genome assemblies are an essential resource for a large variety of evolutionary, ecological, and phylogeographic analyses, as well as for biodiversity monitoring. The presented long-read-based chromosome-level genome assembly of the golden jackal is the first genome assembly for this species and thus provides a solid basis for in-depth genomic studies on the species’ range expansion, population dynamics, adaptability to new environments, and potential as zoonotic disease reservoirs.

## Data Accessibility

All underlying read data and the assembly are available under NCBI GenBank BioProject PRJNA1259920. In addition, the chromosome-level and draft assembly, as well as the annotation results, are available on Zenodo (https://doi.org/10.5281/zenodo.16779415).

## Acknowledgements

We are grateful for funding from the Austrian Partnership Program for Higher Education and Research (APPEAR) project 278 by Austria’s Agency for Education and Internationalisation (OEAD). S.W. acknowledges funding from the Austrian Science Fund (FWF) project I508-B (to P.B.).We thank Anna Kübber-Heiss and Helmut Dier from the Pathology Department of the University of Veterinary Medicine Vienna for access to the frozen tissue collection and the Bioscientia Institut für Medizinische Diagnostik GmbH (Ingelheim, Germany) for providing the PacBio SMRT sequencing service on the PacBio Revio platform.

