## Supplementary Material for "A *de novo* reference genome of the golden jackal, *Canis aureus*"

**Supplementary Material S1. Summary statistics of raw PacBio sequencing data.**

| **PacBio sequencing** |  |
| --- | --- |
| Mean read length | 13,336.4 |
| Mean read quality | 29.7 |
| Median read length | 12,640.0 |
| Median read quality | 30.8 |
| Number of reads | 6,520,091 |
| Read length N50 | 13,319 |
| Total bases | 86,954,770,380 |

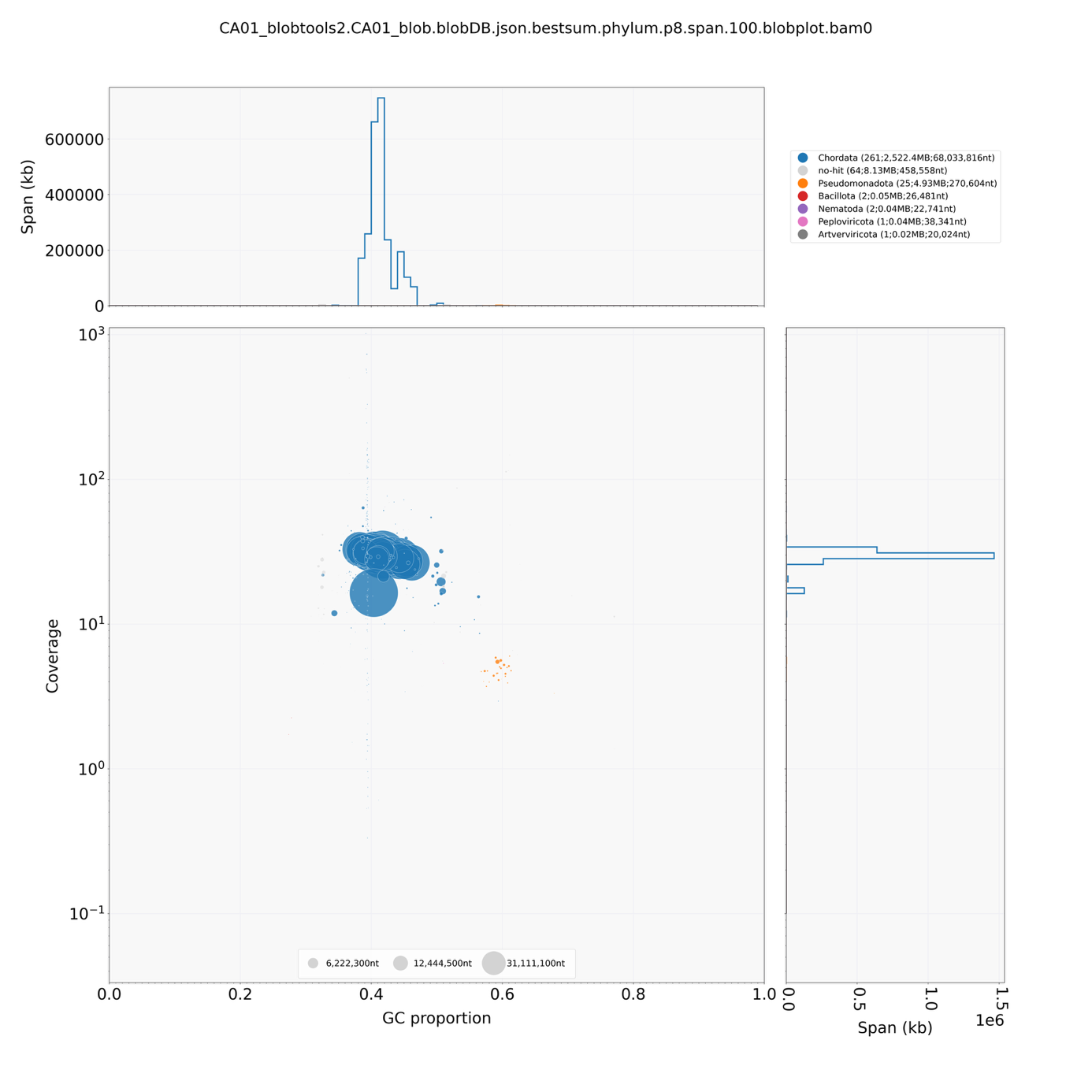

**Supplementary Material S2. BlobPlot of scaffolded golden jackal assembly before removal of contaminated scaffolds.** A cluster of scaffolds with distinct GC-content, coverage, and taxonomic assignment different than the main cluster indicates likely contamination (e.g., orange Pseudomonadota cluster).
